# Treeline ecotones shape the distribution of avian species richness and functional diversity in south temperate mountains

**DOI:** 10.1101/2020.09.23.310177

**Authors:** Tomás A. Altamirano, Devin R. de Zwaan, José Tomás Ibarra, Scott Wilson, Kathy Martin

**Affiliations:** Department of Forest and Conservation Sciences, University of British Columbia, Vancouver, BC, Canada; ECOS (*Ecosystem-Complexity-Society*) Laboratory, Centre for Local Development, Education and Interculturality, Villarrica Campus, Pontificia Universidad Católica de Chile, La Araucanía Region, Chile; Millennium Nucleus Center for the Socioeconomic Impact of Environmental Policies (CESIEP) & Center of Applied Ecology and Sustainability (CAPES), Pontificia Universidad Católica de Chile, Santiago, Chile; Environment and Climate Change Canada, Pacific Wildlife Research Centre, Vancouver, BC, Canada; Department of Biology, 1125 Colonel By Drive, Carleton University, Ottawa ON, K1S 5B6 Canada

**Keywords:** Alpha diversity, Andes, avian community, beta diversity, elevational gradients, environmental filters, high Andean habitats, montane, Patagonia

## Abstract

Mountains produce distinct environmental gradients that may constrain or facilitate both the presence of avian species and/or specific combinations of functional traits. We addressed species richness and functional diversity to understand the relative importance of habitat structure and elevation in shaping avian diversity patterns in the south temperate Andes, Chile. During 2010-2018, we conducted 2,202 point-counts in four mountain habitats (successional montane forest, old-growth montane forest, subalpine, and alpine) from 211 to 1,768 m in elevation and assembled trait data associated with resource use for each species to estimate species richness and functional diversity and turnover. We detected 74 species. Alpine specialists included 16 species (22%) occurring only above treeline with a mean elevational range of 298 m, while bird communities below treeline (78%) occupied a mean elevational range of 1,081 m. Treeline was an inflection line, above which species composition changed by 91% and there was a greater turnover in functional traits (2–3 times greater than communities below treeline). Alpine birds were almost exclusively migratory, inhabiting a restricted elevational range, and breeding in rock cavities. We conclude that elevation and habitat heterogeneity structure avian trait distributions and community composition, with a diverse ecotonal sub-alpine and a distinct alpine community.

## INTRODUCTION

Our understanding of the factors shaping geographic ranges of species is based predominantly on patterns across broad spatial scales ^1^. Fine scale studies across environmental gradients allow us to examine the relationships between range limits, diversity, and environmental features to elucidate the processes by which the environment shapes community assembly ^2^. Mountain ecosystems provide an excellent system for studying these relationships. Comprising nearly a quarter of the global land-base ^3^, mountains produce distinct biotic and abiotic environmental filters over relatively short distances ^4^. For example, with increasing elevation, temperatures decline, the frequency of extreme weather events increase, breeding season durations decrease, and habitat structure becomes more open ^5,6^. Thus, spatially bounded environmental filters shape the distribution and assembly of biological communities across elevational gradients ^7,8^.

Species range limits may be dictated by constraints on total resource availability, and these limits can be visualized as the position of the species in a dynamic and multidimensional ecological niche space ^2,9^. Habitats with greater heterogeneity should facilitate ecological niche divergence, allowing for a greater number of co-occurring species in a given space and structuring the distribution of avian communities across elevational gradients ^10,11^. These effects are pronounced in tropical mountains where habitat specialization among species combined with heterogeneity in vegetation typically generates high diversity turnover (i.e. beta diversity) ^12–15^. For example, species turnover in the wet tropical mountains of Central America is exceptionally high with over a 90% turnover of the avian community within only 500 m of elevation ^14^.

Fewer studies have focused on the influence of heterogeneity on diversity in temperate mountain ecosystems where high seasonality and low species richness should reduce the intensity of inter-specific competition relative to tropical mountains ^13,16,17^. Hypothetically lower levels of competition in temperate mountain ecosystems could enable coexistence among bird species with similar ecological niches ^18^, resulting in large elevational ranges that reflect a broader spectrum of habitat and resource use (i.e. habitat generalists) ^19,20^. If true, we would expect that greater structural heterogeneity within habitats should enhance diversity independent of elevation given that the local community is unsaturated. However, little is known about the relationship between habitat heterogeneity, elevation, and diversity in temperate mountain ecosystems, particularly in the southern hemisphere ^20,21^.

Assessing the covariation of species richness and functional diversity can help identify the mechanisms by which habitat heterogeneity and elevation shape community assembly ^22,23^. Functional diversity is the value, range, and density of behavioral, morphological, and physiological traits (hereafter traits; e.g. breeding strategy, diet) in ecological communities and provides a mechanistic link between organisms and ecosystem function ^23,24^. Functional diversity can explain variation in ecosystem function (e.g. productivity) even if species richness does not ^25^. Differences in how species richness and functional diversity vary across elevational gradients indicate the level of functional redundancy in a community ^26,27^. For instance, communities with high functional redundancy (i.e., support several species with similar trait combinations) can maintain their functional diversity even with reductions in species richness ^23,26^. Conversely, similar changes in species richness and functional diversity across a gradient might indicate low redundancy. Abrupt changes in species and/or traits (functional) along elevational gradients also indicate regions where the environment acts as a strong selective agent in shaping community structure ^28^. Therefore, by addressing how both species and their traits are distributed in mountain ecosystems, we can assess the relative role of habitat heterogeneity and elevation on community identity and function.

We investigated the combined effects of habitat (type and heterogeneity) and elevation on avian diversity and turnover in south temperate mountain ecosystems, using both species richness and functional metrics. Specifically, we assessed: 1) elevational range limits for mountain bird species, 2) patterns of species richness and functional diversity to test whether diversity increases with habitat heterogeneity, and 3) species and functional turnover across elevation and, if any, the relative influence of specific traits supporting this change within and across different mountain habitats. Based on the previously described diversity patterns and linkages with both habitat and elevation, we predicted that in the south temperate Andes: i) elevational range limits of avian communities would be dominated by habitat generalist species (i.e. broad elevational ranges), rather than habitat specialist species with narrow elevational ranges ^14,20^, ii) habitat heterogeneity would further increase species richness and functional diversity within elevations; thus both elevation *per se* and habitat heterogeneity contribute to species richness and functional diversity patterns in the south temperate mountains ^10,21^, and iii) species and functional turnover would be gradual across elevation intervals and habitats, in contrast to the rapid turnover patterns observed in tropical mountains ^14^.

## RESULTS

We recorded 30,969 bird detections of 74 species inhabiting south temperate mountain ecosystems belonging to 26 families and 15 orders (see Appendix S1). Thirty-three bird species (45%) were observed over an elevational range of 1,000 m or more (maximum of 1,524 m, Figure 2). The elevational distributions of birds varied above and below treeline. The avian community above the treeline was comprised of 38 species, 16 of which (22% of all species detected) occurred only in alpine habitats; these habitat specialists had a mean ± SE elevational range of 298 ± 34 m. By contrast, the bird community below treeline included 58 bird species (78% of all species detected) with a mean elevational range of 1,081 ± 58 m. Thus, our prediction that south temperate mountains birds would have wide elevational ranges was supported in the community below treeline but not above treeline. In the lowest section of the elevational gradient, there were seven bird species highly associated with anthropogenic disturbances (excluding two species with only one detection), with a mean elevational range of 426 ± 93 m (Figure 2).

**FIGURE 1.**
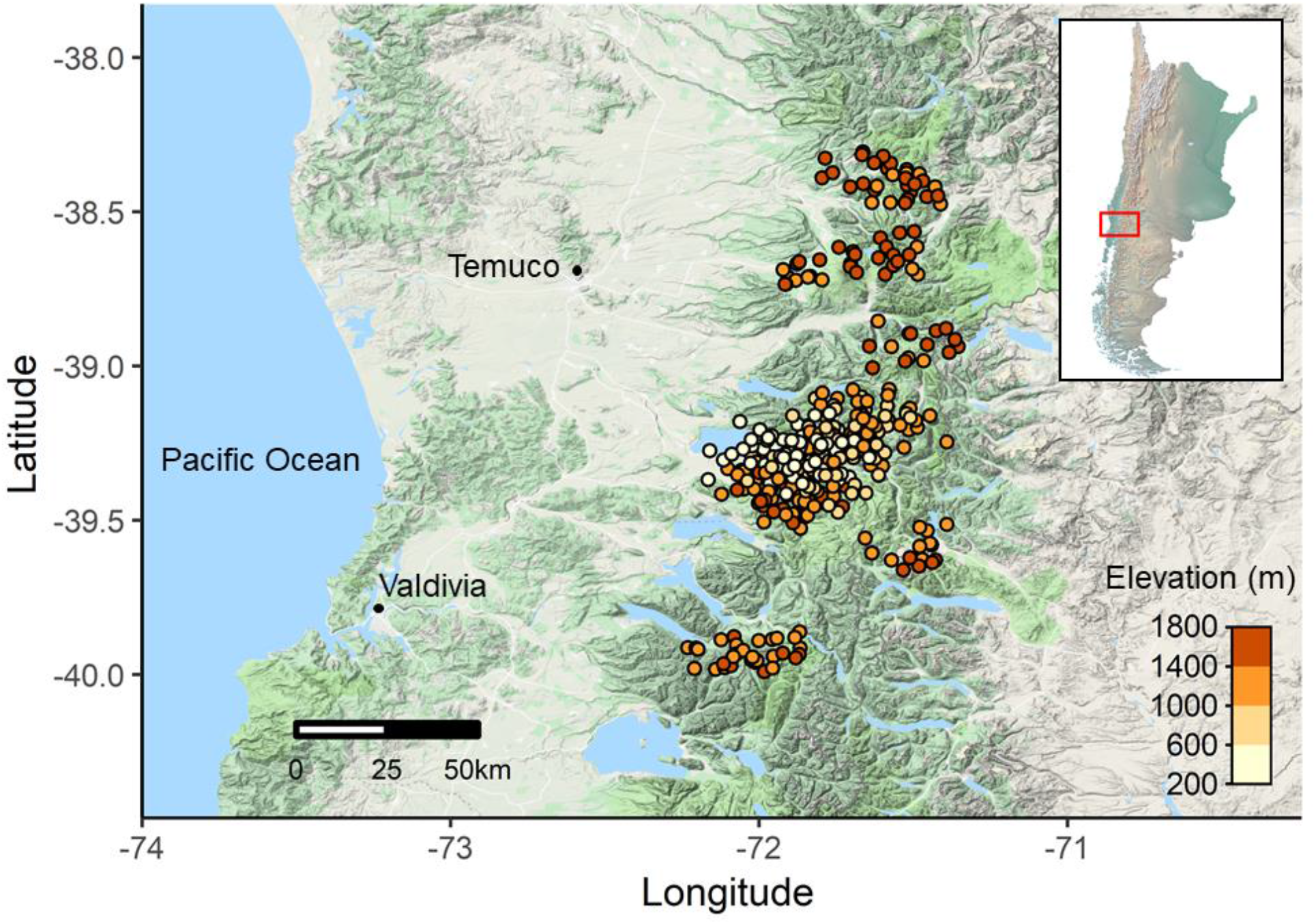
Avian point-count locations conducted between 2010 and 2018 in south temperate Andean mountains, Chile. This map was created by DDZ using R 3.4.4 ^75^ (www.r-project.org/).

**FIGURE 2.**
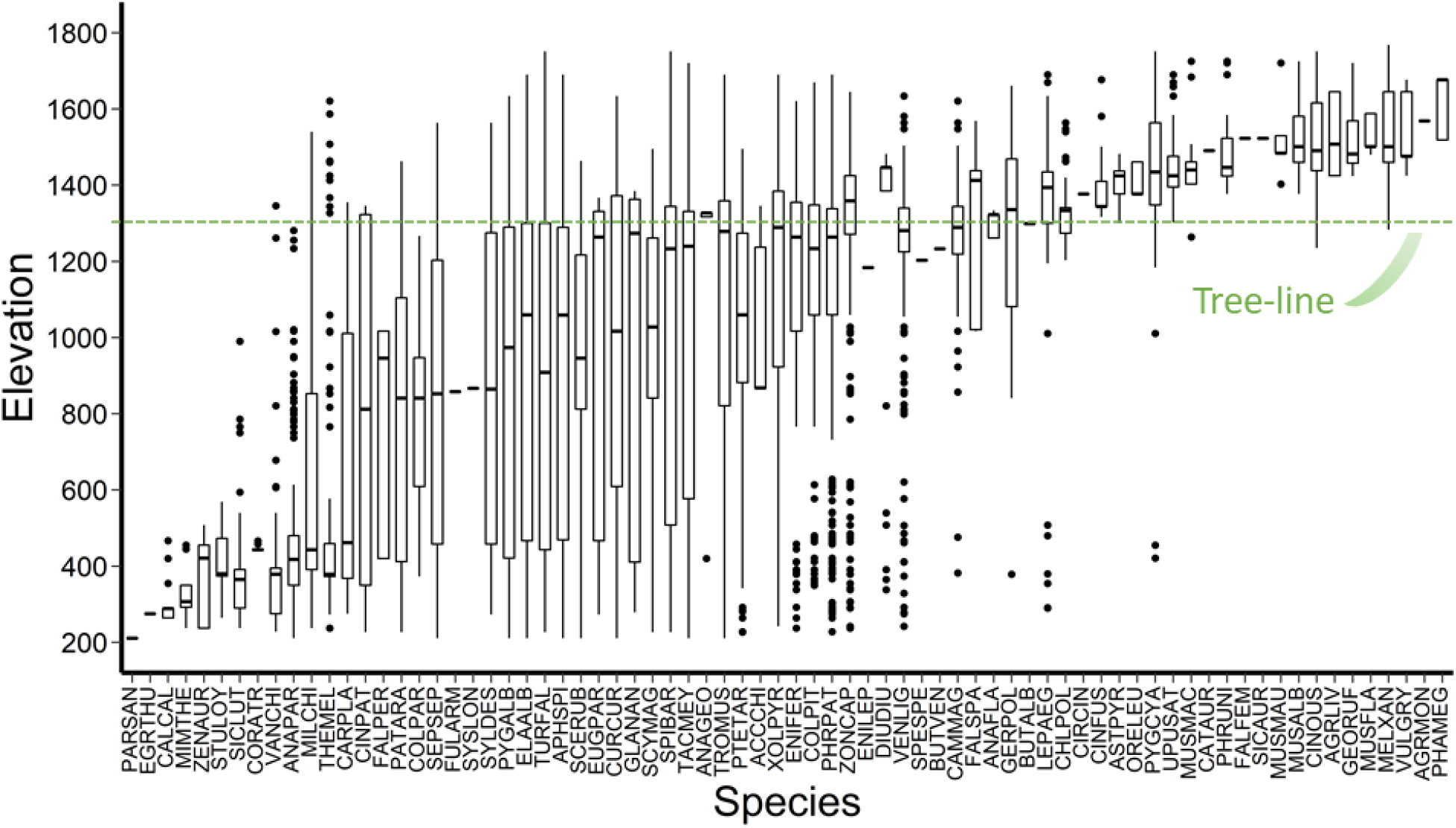
Density weighted box plot of the elevational ranges for 74 bird species inhabiting the south temperate Andes Mountains, Chile. The thick horizontal line within the box represents the median of each species, the boxes represent the first and third quartiles, vertical whiskers represent 1.5 * inter-quartile range (distance between the first and third quartiles), and the outlying points are plotted individually (points that lie outside of the whiskers). The green dashed line represents the average treeline elevation. See species codes in supporting information Appendix S1.

### Bird-habitat relationship across elevations

Variation in species richness was best predicted by models that included habitat type, elevation, and the structural heterogeneity index (Table 3). The combined influence of habitat type and elevation led to a hump-shaped pattern of diversity along elevational gradients, whereby species richness increased until reaching a peak at 1,200–1,400 m asl (old-growth montane forest and subalpine habitats) and then decreased in the alpine (Figure 3a). Since the structural heterogeneity index was included in the top models as an additive term, greater diversity was positively associated with structural heterogeneity independent of either elevation or overall habitat type.

**Table 1.** Functional traits relating to temporal and spatial use of resources for foraging, reproduction and life-history of avian species in Andean temperate mountain ecosystems, south Chile.

| Species name | Diet<sup>a</sup> | Substrate<sup>b</sup> | Nest location<sup>c</sup> | Migratory status<sup>d</sup> | Elevational distribution<sup>e</sup> | Clutch size<sup>f</sup> | Body mass (g)<sup>g</sup> |
| --- | --- | --- | --- | --- | --- | --- | --- |
| Ashy-headed goose (*Chloephaga poliocephala*) | H | G | GO | M | 361 | 5 | 2233.5 |
| Chilean pigeon (*Patagioenas araucana*) | F | F | VO | R | 1236 | 1.5 | 200 |
| Green-backed firecrown (*Sephanoides sephaniodes*) | N | F | VO | M | 1353 | 2 | 5.98 |
| Southern lapwing (*Vanellus chilensis*) | I | G | GO | R | 450 | 3.5 | 323 |
| Black-faced ibis (*Theristicus melanopis*) | I | G | VO | R | 836 | 2.5 | 1250 |
| Variable Hawk (*Geranoaetus polyosoma*) | C | G | VO | M | 820 | 2 | 980 |
| Striped woodpecker (*Veniliornis lignarius*) | I | T | VC | R | 1392 | 3.5 | 39.97 |
| Magellanic woodpecker (*Campephilus magellanicus*) | I | T | VC | R | 764 | 1.5 | 319.5 |
| Chilean flicker (*Colaptes pitius*) | I | T | VC | R | 1320 | 4 | 125 |
| Southern crested caracara (*Caracara plancus*) | C | G | VO | R | 1080 | 3 | 1375 |
| Chimango caracara (*Milvago chimango*) | C | G | VO | R | 1303 | 3 | 291.5 |
| Austral parakeet (*Enicognathus ferrugineus*) | F | F | VC | R | 1384 | 6.5 | 200 |
| Black-throated huet-huet (*Pteroptochos tarnii*) | I | G | VC | R | 1268 | 2 | 144.33 |
| Chucaco tapaculo (*Scelorchilus rubecula*) | I | G | VC | R | 1253 | 2 | 40.35 |
| Magellanic tapaculo (*Scytalopus magellanicus*) | I | G | VC | R | 1268 | 2 | 11.67 |
| Rufous-banded miner (*Geositta rufipennis*) | I | G | RC | M | 297 | 2.5 | 39.5 |
| White-throated treerunner (*Pygarrhichas albogularis*) | I | T | VC | R | 1423 | 3 | 25.6 |
| Patagonian forest earthcreeper (*Upucerthia saturator*) | I | G | GC | M | 388 | 2.5 | 46.25 |
| Buff-winged cinclodes (*Cinclodes fuscus*) | I | G | VC | M | 360 | 3 | 31 |
| Grey-flanked cinclodes (*Cinclodes oustaleti*) | I | G | RC | M | 516 | 3.5 | 26.5 |
| Dark-bellied cinclodes (*Cinclodes patagonicus*) | I | G | GC | M | 1119 | 2.5 | 45.5 |
| Thorn-tailed rayadito (*Aphrastura spinicauda*) | I | T | VC | R | 1479 | 5 | 11.74 |
| Des Murs's wire-tail (*Sylviorthorhynchus desmursii*) | I | F | VO | R | 1291 | 3 | 10.5 |
| Plain-mantled tit-spinetail (*Leptasthenura aegithaloides*) | I | T | VC | R | 679 | 3 | 9.1 |
| Sharp-billed canastero (*Asthenes pyrrholeuca*) | I | F | VO | M | 182 | 3 | 13 |
| White-crested elaenia (*Elaenia albiceps*) | I | F | VO | M | 1479 | 2.5 | 15.62 |
| Tufted tit-tyrant (*Anairetes parulus*) | I | F | VO | R | 779 | 3 | 7.2 |
| Dark-faced ground-tyrant (*Muscisaxicola maclovianus*) | I | G | RC | M | 461 | 2.5 | 23.8 |
| White-browed Ground-tyrant (*Muscisaxicola albilora*) | I | G | RC | M | 348 | 2.5 | 22.6 |
| Fire-eyed diucon (*Xolmis pyrope*) | I | A | VO | R | 1448 | 2.5 | 30.45 |
| Patagonian tyrant (*Colorhamphus parvirostris*) | I | A | VO | M | 894 | 3 | 10.6 |
| Blue-and-white swallow (*Pygochelidon cyanoleuca*) | I | A | RC | M | 740 | 4 | 15.5 |
| Chilean swallow (*Tachycineta leucopyga*) | I | A | VC | M | 1510 | 4 | 16 |
| Southern house wren (*Troglodytes musculus*) | I | F | VC | M | 1479 | 5 | 10.37 |
| Austral thrush (*Turdus falcklandii*) | F | G | VC | R | 1524 | 3 | 78.75 |
| Grassland yellow-finch (*Sicalis luteola*) | G | G | GO | R | 303 | 4 | 16.25 |
| Patagonian sierra-finch (*Phrygilus patagonicus*) | G | G | VO | M | 1462 | 3.5 | 21.3 |
| Plumbeous sierra-finch (*Phrygilus unicolor*) | G | G | RC | M | 348 | 2.5 | 21.9 |
| Yellow-bridled finch (*Melanodera xanthogramma*) | F | G | RC | M | 485 | 3 | 36 |
| Common diuca-finch (*Diuca diuca*) | G | G | VO | R | 299 | 3 | 33 |
| Rufous-collared Sparrow (*Zonotrichia capensis*) | G | G | GO | M | 1408 | 4 | 23.5 |
| Austral black bird (*Curaeus curaeus*) | I | G | VO | R | 1423 | 4.5 | 90 |
| Long-tailed meadowlark (*Sturnella loyca*) | I | G | GO | R | 305 | 4 | 104.75 |
| Black-chinned siskin (*Spinus barbatus*) | G | G | VO | M | 1524 | 4.5 | 15.83 |

<sup>a</sup> I = insectivorous, C = carnivorous, G = granivorous, N = nectarivores, F = frugivorous, H = herbivorous <sup>76,77</sup>. The primary diet is reported and it was used for analysis.
<sup>b</sup> G = ground, A = air, F = foliage, T = timber <sup>76,77</sup> (complemented with our own field observations). The primary foraging substrate is reported and it was used for analysis.
<sup>c</sup> VC = vegetation and cavity-nester, VO = vegetation and open cup nester, GC = ground and cavity-nester, GO = ground and open cup nester, RC = rock and cavity-nester, RO = rock and open cup nester <sup>34,77,78</sup>.
<sup>d</sup> M = migrant, R = resident <sup>30,76,77</sup>.
<sup>e</sup> This study.
<sup>f</sup> 77–82.
<sup>g</sup> <sup>81,83</sup> complemented with field data (Altamirano, Unpublished data).

**Table 2.** Habitat attributes, species richness, and functional diversity per elevational interval and distance to the treeline (i.e. elevation of the point-count minus the elevation of the closest treeline) in south temperate mountains, Chile. Positive and negative intervals are above and below the treeline, respectively. The elevational intervals are arranged from highest to lowest elevation. Each metric is presented as mean (SE).

| Distance to treeline | n<sup>a</sup> | Habitat types<sup>b</sup> | Structural heterogeneity<sup>c</sup> | Species richness<sup>d</sup> | Functional richness<sup>e</sup> | Functional dispersion | CWM elevational distribution<sup>f</sup> |
| --- | --- | --- | --- | --- | --- | --- | --- |
| (+) 200–299 | 36 | AL | 2.36 (0.10) | 2.48 (0.11) | 0.04 (0.01) | 0.09 (0.02) | 552.27 (42.28) |
| (+) 100–199 | 156 | AL | 2.54 (0.05) | 2.62 (0.08) | 0.06 (0.01) | 0.13 (0.01) | 706.96 (26.44) |
| (+) 0–99 | 234 | AL, SA | 3.33 (0.06) | 5.01 (0.18) | 0.12 (0.00) | 0.20 (0.01) | 1110.22 (26.42) |
| (-) 0–99 | 394 | SA, OM | 3.98 (0.03) | 7.82 (0.04) | 0.13 (0.00) | 0.26 (0.00) | 1421.33 (4.01) |
| (-) 100–199 | 296 | SA, OM | 4.04 (0.02) | 7.59 (0.05) | 0.11 (0.00) | 0.26 (0.00) | 1434.58 (3.02) |
| (-) 200–299 | 36 | OM | 4.00 | 6.73 (0.11) | 0.09 (0.01) | 0.26 (0.00) | 1455.43 (5.47) |
| (-) 300–399 | 152 | OM, SM | 3.75 (0.05) | 7.25 (0.07) | 0.10 (0.00) | 0.25 (0.00) | 1417.96 (3.71) |
| (-) 400–499 | 130 | OM, SM | 3.64 (0.04) | 6.39 (0.07) | 0.07 (0.00) | 0.26 (0.00) | 1407.80 (3.82) |
| (-) 500–599 | 222 | OM, SM | 3.87 (0.02) | 6.70 (0.06) | 0.08 (0.00) | 0.25 (0.00) | 1410.16 (2.91) |
| (-) 600–699 | 23 | OM, SM | 3.65 (0.10) | 5.87 (0.13) | 0.08 (0.01) | 0.25 (0.01) | 1397.92 (13.11) |
| (-) 700–799 | 18 | SM | 3.67 (0.11) | 6.10 (0.16) | 0.10 (0.01) | 0.24 (0.02) | 1448.15 (10.63) |
| (-) 800–899 | 29 | SM | 3.48 (0.11) | 6.03 (0.08) | 0.07 (0.01) | 0.25 (0.01) | 1391.57 (25.02) |
| (-) 900–999 | 277 | SM | 3.81 (0.02) | 6.26 (0.04) | 0.09 (0.00) | 0.25 (0.00) | 1423.35 (3.74) |
| (-) 1000–1099 | 130 | SM | 3.78 (0.04) | 5.77 (0.07) | 0.08 (0.00) | 0.25 (0.00) | 1412.03 (9.15) |
| (-) 1100–1199 | 69 | SM | 3.91 (0.03) | 5.63 (0.06) | 0.07 (0.01) | 0.24 (0.01) | 1414.80 (13.75) |

<sup>a</sup> Sample size for each elevational interval.
<sup>b</sup> AL: Alpine, SA: Subalpine, OM: Old-growth montane forest, SM: Successional montane forest.
<sup>c</sup> Additive index of the structural habitat attributes (i.e. tree canopy, dead trees, understory, shrub, snow, tundra, and rock).
<sup>d</sup> Predicted species richness from the top model in Table 3.
<sup>e</sup> Higher values indicate a greater volume of the potential niche space is occupied.
<sup>f</sup> Community-weighted mean of the elevational distribution trait.

**Table 3.**
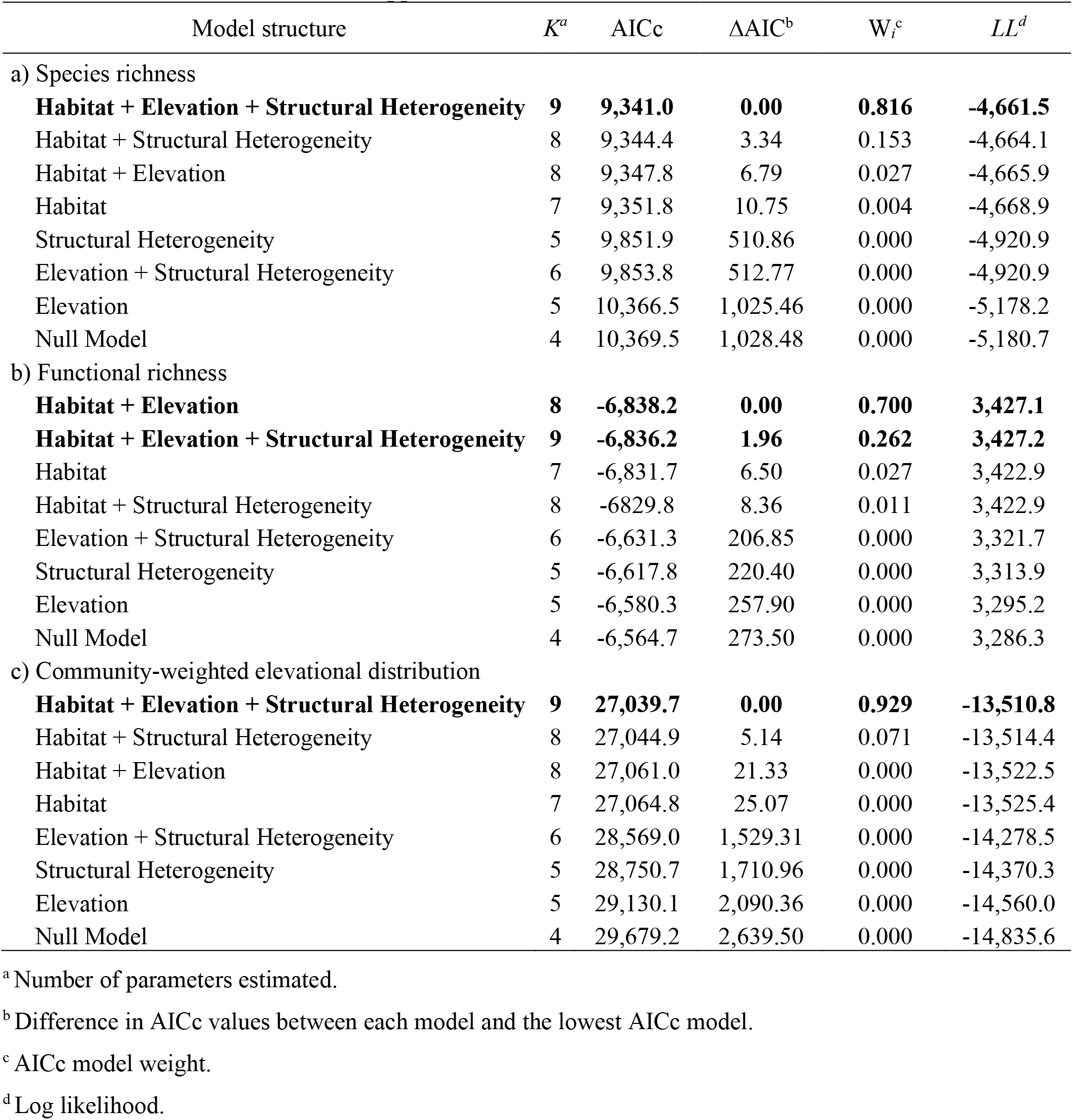
Model rankings for species richness and functional diversity in relation to habitat type, elevation, and structural heterogeneity (i.e. additive index of the structural habitat attributes) for avian surveys conducted in south temperate mountains, Chile. Year and site were random terms in all models. Bold indicates best-supported models.

| Model structure | $K^a$ | AICc | $\Delta AIC^b$ | $W_i^c$ | $LL^d$ |
| --- | --- | --- | --- | --- | --- |
| a) Species richness |  |  |  |  |  |
| **Habitat + Elevation + Structural Heterogeneity** | **9** | **9,341.0** | **0.00** | **0.816** | **-4,661.5** |
| Habitat + Structural Heterogeneity | 8 | 9,344.4 | 3.34 | 0.153 | -4,664.1 |
| Habitat + Elevation | 8 | 9,347.8 | 6.79 | 0.027 | -4,665.9 |
| Habitat | 7 | 9,351.8 | 10.75 | 0.004 | -4,668.9 |
| Structural Heterogeneity | 5 | 9,851.9 | 510.86 | 0.000 | -4,920.9 |
| Elevation + Structural Heterogeneity | 6 | 9,853.8 | 512.77 | 0.000 | -4,920.9 |
| Elevation | 5 | 10,366.5 | 1,025.46 | 0.000 | -5,178.2 |
| Null Model | 4 | 10,369.5 | 1,028.48 | 0.000 | -5,180.7 |
| b) Functional richness |  |  |  |  |  |
| **Habitat + Elevation** | **8** | **-6,838.2** | **0.00** | **0.700** | **3,427.1** |
| **Habitat + Elevation + Structural Heterogeneity** | **9** | **-6,836.2** | **1.96** | **0.262** | **3,427.2** |
| Habitat | 7 | -6,831.7 | 6.50 | 0.027 | 3,422.9 |
| Habitat + Structural Heterogeneity | 8 | -6,829.8 | 8.36 | 0.011 | 3,422.9 |
| Elevation + Structural Heterogeneity | 6 | -6,631.3 | 206.85 | 0.000 | 3,321.7 |
| Structural Heterogeneity | 5 | -6,617.8 | 220.40 | 0.000 | 3,313.9 |
| Elevation | 5 | -6,580.3 | 257.90 | 0.000 | 3,295.2 |
| Null Model | 4 | -6,564.7 | 273.50 | 0.000 | 3,286.3 |
| c) Community-weighted elevational distribution |  |  |  |  |  |
| **Habitat + Elevation + Structural Heterogeneity** | **9** | **27,039.7** | **0.00** | **0.929** | **-13,510.8** |
| Habitat + Structural Heterogeneity | 8 | 27,044.9 | 5.14 | 0.071 | -13,514.4 |
| Habitat + Elevation | 8 | 27,061.0 | 21.33 | 0.000 | -13,522.5 |
| Habitat | 7 | 27,064.8 | 25.07 | 0.000 | -13,525.4 |
| Elevation + Structural Heterogeneity | 6 | 28,569.0 | 1,529.31 | 0.000 | -14,278.5 |
| Structural Heterogeneity | 5 | 28,750.7 | 1,710.96 | 0.000 | -14,370.3 |
| Elevation | 5 | 29,130.1 | 2,090.36 | 0.000 | -14,560.0 |
| Null Model | 4 | 29,679.2 | 2,639.50 | 0.000 | -14,835.6 |

<sup>a</sup> Number of parameters estimated.
<sup>b</sup> Difference in AICc values between each model and the lowest AICc model.
<sup>c</sup> AICc model weight.
<sup>d</sup> Log likelihood.

**FIGURE 3.**
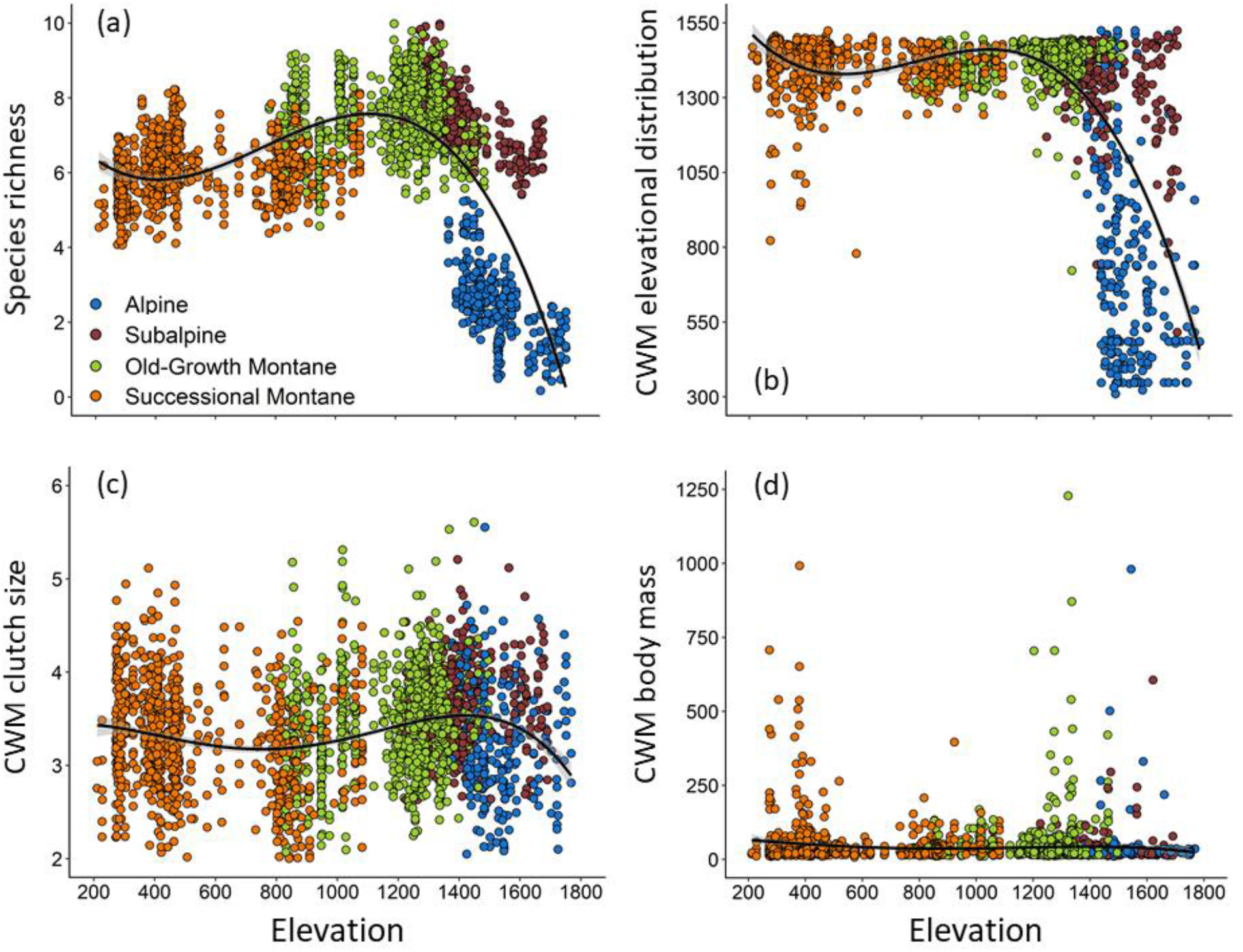
Combined effect of elevation and habitat type on species richness and community-weighted mean (CWM) of functional traits in the south temperate Andes, Chile. (a) Predicted values of species richness (based on the top model in Table 3), (b) CWM of elevational distribution, (c) CWM of clutch size, and (d) CWM of body mass. Every point represents a point-count survey carried out in successional montane forest (>50% canopy cover, 35-100 years old), old-growth montane forest (>50% canopy cover, >200 years old), subalpine (5-50% canopy cover), or alpine (<5% canopy cover).

Habitat type, elevation, and the structural heterogeneity index also best explained variation in functional diversity in the Andean temperate mountains (Table 3). Functional richness varied significantly across habitats, with the highest complement of species exhibiting different traits in subalpine habitats, followed by old-growth montane forests. In contrast, the lowest functional richness values were found in alpine habitats followed by successional montane forests (Table 2). Consequently, we found a strong and positive linear association between species and functional richness (r^2^ = 0.76), indicating low redundancy in avian communities inhabiting the south temperate Andes. The community-weighted mean of elevational distribution was best explained by a model including habitat type, elevation, and the structural heterogeneity index (Figure 3b, Table 3), with relatively stable values across elevation below the treeline, after which the elevational distribution of the community decreased. In contrast, the community-weighted means of clutch size and body mass did not vary with elevation (Figure 3c, d).

### Species and functional turnover across elevational gradients

The cluster analysis showed a relatively low species turnover across elevation below treeline, with an average 28% turnover in species composition among 100 m elevational intervals (range of 12–46%). There was a clear inflection line in the composition of the bird community above the treeline, where the community changed by 91% compared to the community below the treeline (range of 82–96%, Figure 4).

**FIGURE 4.**
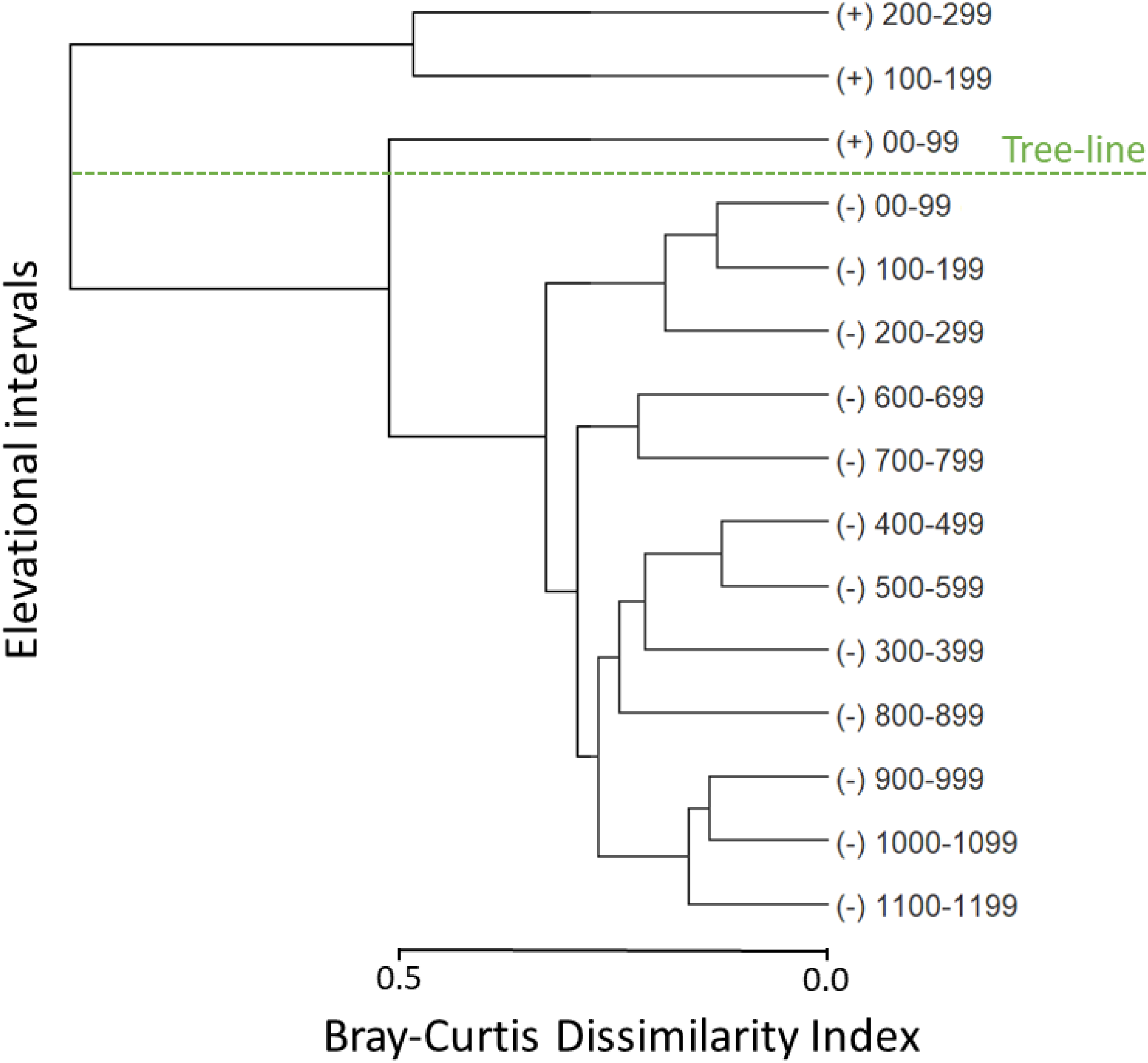
Density-weighted bird species cluster across 100-m elevational intervals based on distance to the closest treeline, for each point-count, in the south temperate Andes Mountains, Chile. Positive and negative intervals are above and below the treeline, respectively. A Bray-Curtis dissimilarity index of 0 means a complete overlap in species, while 1 indicates no shared species in the bird community.

With respect to functional turnover, functional dispersion (i.e. functional similarity among species in a community) varied significantly among bird communities across different habitats. Alpine bird communities had the lowest dispersion (more similarity) compared to the bird communities below treeline (F_Disp_ = 0.12 ± 0.007), while habitats at mid elevation (i.e. old-growth montane forests, F_Disp_ = 0.26 ± 0.001; subalpine, F_Disp_ = 0.26 ± 0.002) had the highest dispersion (less similarity). Larger functional distances between centroids were found in pairwise comparisons (i.e. habitat*_i_*/habitat*_j_*) that included alpine bird communities (effect size ± 95% confidence intervals ranged from 0.33 ± 0.01 to 0.43 ± 0.01). Thus, the turnover was between 2.2 and 3 times higher when comparing alpine bird communities to those below treeline than for comparisons among bird communities below the treeline (subalpine/old-growth montane forest: 0.17 ± 0.00; subalpine/successional montane forest: 0.19 ± 0.00; old-growth montane forest/successional montane forest: 0.17 ± 0.00). The most influential traits driving functional turnover were migratory status, elevational distribution, and breeding strategy (Figure 5). Alpine bird communities were distinct from those below the treeline as they were mainly migratory, inhabited a restricted elevational range, and bred in rock cavities. Thus, our prediction of a gradual turnover with elevation is supported only below the treeline. Above treeline, we found a strong species and functional turnover in the community.

**FIGURE 5.**
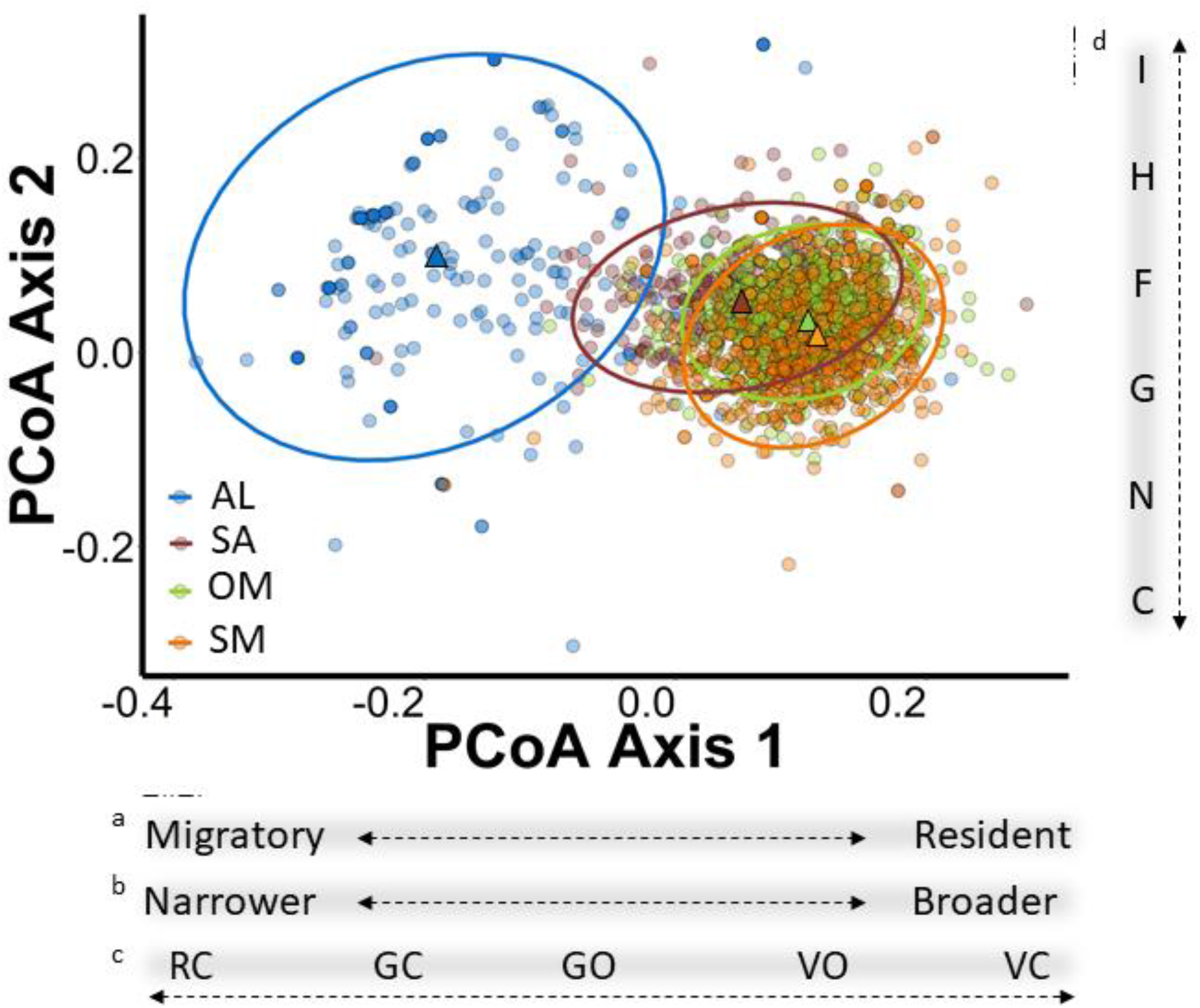
Functional distance of mountain avian bird communities in south temperate Andes, Chile. Functional position (circles) and centroids (triangles) of each mountain bird community associated with habitat type, ellipses represent the 95% confidence level. The relationship between PCoA Axes (Principal Coordinate Analysis) and the most influential functional traits are indicated in the panels below the figure for PCoA Axis 1: a. Migratory status, b. Elevational distribution, c. Nest location, and in the vertical panel to right of figure for PCoA Axis 2: d. Diet. AL: Alpine, SA: Subalpine, OM: Old-growth montane forest, SM: Successional montane forest (see Table 1 for codes and categories of functional traits).

## DISCUSSION

In south temperate Andean mountains, we demonstrated that most mountain bird species are distributed across wide elevational ranges, from valley bottom to treeline. Treeline represents an ‘inflection line’ in avian communities across the broader elevational gradient, above which species richness and functional diversity exhibit the greatest turnover, resulting in a highly specialized alpine bird community. Below treeline, elevational range limits were quite broad, with 65% of species found from lowland to sub-alpine forests and 45% exhibiting elevational ranges of 1,000 to 1,524 m. This contrasts strongly with expectations for tropical mountain birds which often have distributions restricted to elevational intervals of 500 m or less ^13,14,29^, but supports previous findings for forest-dwelling species in the south temperate Andes ^30,31^. In contrast, above treeline, species had more compressed elevational distributions (~300 m), with 16 of the 38 species observed in the alpine occurring exclusively in this habitat. Thus, our results highlight the difference in distributions and habitat specialization between species above and below treeline.

Species richness was strongly structured by habitat across elevation ^21,32^. The greatest diversity was observed in the ecotonal subalpine habitats which were associated with the highest habitat heterogeneity and supported a broad spectrum of ecological bird requirements: forest generalists, old-growth montane forest specialists, ecotone specialists (e.g. Patagonian Forest Earthcreeper, *Upucerthia saturatior*), and the occasional alpine species that forages within the subalpine-alpine transition zone (e.g. Plumbeous sierra-finch, *Phrygilus unicolor*). While most species observed in forested mountain habitats occupied the entire elevational range below the treeline (habitat generalists) ^20^, greater structural heterogeneity in old-growth montane forests and the subalpine, as well as within habitats, was associated with higher species richness. Thus, more heterogenous vegetation at multiple spatial scales can support greater diversity and abundance in avian communities ^33^, likely by offering more diverse nesting resources and broader niche space ^34^. Accelerating land-use at lower elevation may also push some forest specialists upslope, potentially inflating the diversity peaks observed at higher elevation. For example, forest specialists such as the Magellanic Woodpecker (*Campephilus magellanicus*) and Austral Parakeet (*Enicognathus ferrugineus*) had lower elevational limits of approximately 800 m, likely reflecting the loss of their preferred old-growth forests at lower elevations ^35,36^. These findings suggest greater productivity in high elevation forested habitats compared to other mountain habitats and highlight the importance of conserving old-growth montane forests in the southern Andes ^37,38^.

Mountain habitats with high structural heterogeneity were also associated with the greatest overlap of functional trait distributions. Addressing the distribution of traits across elevation provides a more comprehensive assessment of the resiliency of a community and therefore can be more informative than species richness ^36,39,40^. We found a non-saturating relationship between species and functional richness, which is expected in communities with low functional redundancy (i.e. few organisms resembling each other in their traits) ^41^. Both measures of richness increased with elevation up to treeline and then decreased above treeline, indicating low redundancy in the whole elevational gradient in the south temperate Andes ^10^. Therefore, relative to a more diverse community composed of multiple species with similar ecological roles, such as tropical mountain bird communities in Central America ^42^, the addition or loss of a species would have a proportionately greater influence on the resiliency of south temperate mountain bird communities ^43^. We acknowledge that there are other mountain bird species with lower densities (1% of the bird detections) that might add some functional traits, although these are likely evenly distributed across elevations as these species were mainly raptors or wetland birds with broader elevational ranges that occur infrequently and do not necessarily rely on mountain habitats to breed or forage.

Species richness decreased above the treeline, but habitat specialization within the bird community increased. The magnitude of the species turnover at treeline is similar to that reported across elevations for tropical mountains ^35,44^. This compositional change likely reflects a combination of multiple habitat and climate filters. The absence of some habitat structures, such as trees (alive and dead), is a strong factor supporting communities of cavity-nesting species (57% of species below treeline nested in tree cavities) ^34^. Although species occupying over a 1000 m gradient below treeline would experience a very wide gradient of climatic conditions, especially early in the breeding season, harsher climatic conditions above treeline (more persistent snow and colder temperatures with more storms) may further promote turnover, selecting for species that are well adapted to surviving and breeding in extreme and exposed environments ^6,13^.

We also demonstrated support for functional turnover, above treeline only, as the alpine bird community showed both significantly less functional dispersion compared to bird communities at or below treeline, and the most distant centroid in multi-dimensional trait space ^45^. The latter indicates specialization as alpine birds are functionally closer to each other than bird communities in other mountain habitat types. Functional turnover was mainly driven by seasonal use of mountain habitats, elevational distribution in mountains, and breeding strategy. Alpine bird communities occupied a unique position in trait space, consisting of mainly migratory species restricted to relatively small elevational intervals that predominantly breed in natural rock cavities. This contrasted with communities below treeline, where functional traits like seasonal habitat use (migration) and breeding strategy were more diverse and broadly distributed across elevation. Strong seasonality in temperate mountains may select for broader physiological thermal tolerances compared to the tropical Andean mountains ^16^. However, thermal tolerance of the alpine bird community might be narrowed by migration behavior that allows for specialization within narrow elevational bands without having to cope with strong seasonal changes in temperature ^46^. Additionally, the prevalence of rock cavity nesting strategies in the alpine may be in response to a higher probability of desiccation in drier alpine environments with little vegetation cover and/or higher predation risk for open cup ground nests ^47^. Regardless of the mechanism, the propensity to nest in rock cavities is a key trait that distinguishes the alpine from communities below treeline in south temperate Andean ecosystems.

## Conclusions

By addressing species richness and functional diversity across elevational gradients, we demonstrate peaks in avian diversity at mid-to-high elevations (i.e. old-growth montane forest and subalpine) in the south temperate Andes. Mountain habitats characterized by high structural heterogeneity were associated with the greatest overlap of species and functional trait distributions. Importantly, structural heterogeneity within habitats was also positively associated with diversity, indicating the critical role of diverse vegetative structure in promoting productive communities. We therefore stress the importance of considering habitat structure and functional traits when assessing theories of biogeography. We also highlight the unique value of temperate high elevation habitats with a diverse ecotonal sub-alpine, as well as, a taxonomically and functionally distinct alpine avian community. Future research directly addressing additional niche dimensions (e.g. physiological tolerances, life-history variation) and incorporating evolutionary histories through phylogenetic analyses, would further our understanding of community assembly for mountain avifauna. An increasingly variable climate and rapid land-use change is threatening mountain biodiversity by compressing realized elevational ranges ^7,48–50^. Species richness and functional diversity can inform habitat protection and management strategies to promote vegetative heterogeneity and improve the persistence of mountain bird diversity ^51–53^.

## METHODS

### Study area

We investigated avian diversity across elevational gradients from 211 to 1,768 m of elevation in the south temperate Andes in La Araucanía and Los Ríos Regions, Chile (38–40°S latitude, north-south distance of 182 km; Figure 1). The average treeline elevation is 1,300 m above sea level (asl), and permanent snow and/or rock terrains start around 1,800 m elevation. Vegetation structure varies across elevation within- and among-mountains based on the timing of natural disturbances (e.g. volcanic eruptions) and/or land use change over time (e.g. agroforestry). We identified four habitat types outlined in Nagy & Grabherr ^4^ and Boyle & Martin ^54^ which are differentiated by elevation and habitat structure: (a) successional montane forests (200–1000 m asl, > 50% tree cover, 35-100 years old); (b) old-growth montane forests, between 800 m asl and the forest-line (i.e., the end of continuous forest; > 50% tree cover, > 200 years old); (c) subalpine, an ecotonal habitat with a mix of highland herbaceous meadows, shrubs, and sparse patches of trees and/or krummholz existing between the forest-line and the treeline (i.e. the highest elevation with trees ~ 3 m in height; 5–50% tree cover); (d) alpine, high Andean tundra habitats occurring above the treeline and characterized by perennial herbaceous plants, shrubs, few or no trees, and a strong influence of volcanic disturbances (< 5% tree cover).

### Avian surveys

From 2010 to 2018, over nine austral breeding seasons (October to February), we conducted 2,202 point-transect surveys ^55^ (Table 2). We systematically established point-count stations across elevational gradients with a minimum distance of 125 m between stations ^41^. Each point-count survey lasted six minutes and was conducted between 0515 and 1000 hours. We recorded number of individuals for every diurnal bird species identified by sight and sound ^56^, and estimated the distance to all detected birds within two concentric bands of 25 m (i.e. 0-25 m, 26-50 m) ^41^.

### Abiotic and biotic covariates

We recorded the elevation of each point-count station using a Global Positioning System (GPS, Garmin). Within 50 m of each point-count station, we estimated the percent cover of the following habitat structural components: tree canopy, dead trees, understory, shrub, snow, tundra, and rock. Using the percent cover estimations, we calculated the habitat ‘structural heterogeneity index’ by summing the percent abundance (0 to 1) of each structural component ^32,33^, which often totaled greater than 1 due to the vertical overlap among structural layers (e.g. canopy and understory covers). This index is independent of the type of structure, as it is based on the diversity of structures, and thus, it is possible to have the same structural heterogeneity index value above and below treeline. We estimated cloud cover (categorically classified into eighths, where 0 = no cloud and 1 = 100% cloud cover) and we recorded temperature (°C), wind speed (m/s), and relative humidity (%) during each survey using a hand-held weather station (Kestrel-meters 3000/3500/4000, Birmingham, MI).

### Species functional traits

To analyze functional diversity across elevation, we built a multidimensional ecological niche for each avian community (bird species detected in a point count) by classifying each species using seven discrete and continuous traits ^10,57^ (Table 1). Categorical traits included diet, foraging substrate, nest location and type, and migratory status. Clutch size was ordinal, while body mass and elevational distribution were continuous traits. These traits were selected because they can be directly related with spatial and temporal resource use (type and quantity) and ecosystem function across elevational gradients. Diet expresses trophic linkages in ecological networks and can be associated with environmental productivity, as well as the extent of ecological processes such as predation, seed dispersion, and pollination ^58,59^. Foraging substrate reflects habitat associations, while nest location and type indicates breeding site selection and different species interactions (e.g. commensal networks) ^34^. Migratory status accounts for temporal dynamics of bird communities informing whether birds use mountains across the complete annual cycle or just during the breeding season ^60^, while elevational distribution is a spatial metric of resource use within mountain habitats. Clutch size characterizes the potential fecundity for a community and the diversity of breeding strategies ^61^, and body mass is related to thermal and energetic constraints and habitat heterogeneity ^62,63^.

### Analysis

#### Probability of species detection and bird densities

We had sufficient data to estimate density for 44 species (species detected in ≥ 10 different point-count surveys), which accounted for 99% of the total bird detections. We analyzed point-count data using Multinomial Poisson Mixture Models in a Multiple-Covariate Distance Sampling framework ^64^, which allowed us to correct the estimated densities by the probability of detection of each species based on distance and other spatiotemporal covariates ^65^. To estimate detection and density of each species across elevation we used maximum-likelihood methods in R-Unmarked ^66^. For details, see protocol used by Ibarra and Martin ^41^.

#### Avian diversity distributions

We estimated species richness using Linear Mixed-Effect Models (LMMs) with a normal distribution and we used the Akaike’s Information Criterion (AIC) approach to select the best fit models ^67^. Elevation, habitat type, and the structural heterogeneity index were included in the models as fixed effects. Elevation and habitat type were moderately correlated (−0.59). Therefore, to account for the elevation within each habitat we took the residuals from an elevation by habitat regression and included this as a covariate when habitat was also in the model ^68^. Year and site were included as random effects, allowing us to include bird richness and abundances from the same year and site while taking into account any inherent capacity of each year and/or site to have lower or higher numbers of birds ^68^. Model support was assessed using model weights and the AICc value (i.e. weight > 0.8 and AICc < 2.0 were considered the best-supported models) ^69^.

To assess functional diversity across elevational gradients, we estimated two metrics per point-count: functional richness and community-weighted mean. Functional richness represents the ecological niche volume filled by species in a community (non-density weighted metric) ^70^ and was calculated using species traits and the observed bird species richness per point-count ^36^. Community-weighted mean, or the average values of specific traits in a community, was calculated for the values of continuous traits (i.e. elevational distribution, clutch size, and body mass) in combination with the estimated species densities per point-count ^10^.

#### Avian diversity turnover

To calculate the species turnover across elevation, we divided the elevational gradient into 15 intervals of 100 m each ^14,32,58^. To standardize the elevation of the habitat types across mountains, we calculated the elevational distance to the closest treeline for each point-count (positive numbers refer to a distance above the treeline while negative numbers are below the treeline). Species turnover was calculated using the density weighted Bray-Curtis dissimilarity index, calculated as:

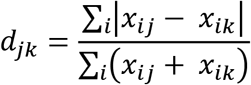

Where *x_ij_* and *x_ik_* are the number of individuals for species *i* and elevational intervals *j* and *k* ^71^. The Bray-Curtis dissimilarity index is bounded between 0 and 1, where 0 means that two given elevation intervals have the same species composition and 1 indicates no species overlap.

To examine functional turnover, we calculated functional dispersion and distance of the traits in bird communities ^72^. Functional dispersion is the mean distance of individual species from the centroid of all species in a community and was calculated by combining species traits with the estimated species density per point-count survey ^73^. Functional distance, the Euclidean distance between the non-density weighted centroids of two communities in a trait space, was calculated using a presence/absence matrix and a Gower distance trait matrix ^28^. From the resulting 2.3 million pairwise comparisons, we assessed the effect of mountain habitat types on the functional distances between the centroids using the following mixed-effect model structure:

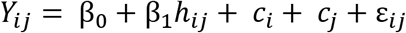

Where *Y_ij_* represents the functional distance between the centroids of communities *i* and *j*, the term *h_ij_* is the fixed effect of habitat pairwise comparisons (habitat*_i_*/habitat*_j_*), both *c_i_* and *c_j_* are random effects terms representing the identity of each community (point-count survey) to account for the non-independence multiple pairwise comparisons with a given bird community (i.e. values in distance matrices), and ε*_ij_* is the independent error term ^45^.

We used R package ‘FD’ to conduct all functional diversity analyses ^73,74^. We conducted Linear Mixed-effect Models to assess the effects of elevation, habitat, and structural heterogeneity index in the same manner that we performed for species richness (see above). Furthermore, we assessed differences in functional dispersion among habitats using one-way ANOVAs. All analyses were conducted using R 3.4.4 ^75^.

## ACKNOWLEDGMENTS

We are grateful for the strong support from Jerry Laker, A. Dittborn, R. Timmerman, M. Sabugal, and C. Délano. Thanks to A. Vermehren, Fernando Novoa, Camila Bravo, Anneka Vanderpas, and Kristina Hick for their support in the field and lab work.

## Permits

Support and permissions to investigate in public protected areas were given by The Chilean Forestry Service (11/2009 IX, 13/2015 IX, RNMCH 892127/2018), and in private protected areas were given by El Cañi Nature Sanctuary, Kawelluco Private Sanctuary, The Huilo Huilo Biological Reserve, Kodkod: Lugar de Encuentros.

## Author contributions

TAA, DRD, JTI, and KM conceived the idea, with input from SW; all authors collected the data; TAA carried out the analyses with support of DRD and JTI. The manuscript was written by TAA. All authors made substantial contributions to the interpretations of results and editing the manuscript.

