## Supplementary material for "Treeline ecotones shape the distribution of avian species richness and functional diversity in south temperate mountains"

### Supporting information

**Appendix S1** Bird species detected in all point count surveys in four mountain habitats in south temperate Andes, Chile (n=74 species). Asterisks show species with enough data for our detectability-density estimation models.

| Species name | Species code | Habitat[Δ] | | | |
|---|---|---|---|---|---|
| | | AL | SA | OM | SM |
| Ashy-headed goose (*Chloephaga poliocephala*)* | CHLPOL | 1 | 1 | 1 | |
| Spectacled Duck (Speculanas specularis) | SPESPE | | | 1 | |
| Yellow-billed pintail *(Anas georgica)* | ANAGEO | | | 1 | 1 |
| Yellow-billed teal (*Anas flavirostris)* | ANAFLA | | | 1 | |
| California quail *(Callipepla californica)* | CALCAL | | | | 1 |
| Chilean pigeon (*Patagioenas araucana*)* | PATARA | | 1 | 1 | 1 |
| Eared dove *(Zenaida auriculata)* | ZENAUR | | | | 1 |
| Band-winged nightjar *(Systellura longirostris)* | SYLSON | | | 1 | |
| Green-backed firecrown (*Sephanoides sephaniodes*)* | SEPSEP | 1 | 1 | 1 | 1 |
| White-sided hillstar *(Oreotrochilus leucopleurus)* | ORELEU | 1 | | | |
| Plumbeous rail *(Pardirallus sanguinolentus)* | PARSAN | | | | 1 |
| Red-gartered coot *(Fulica armillata)* | FULARM | | | | 1 |
| Southern lapwing (*Vanellus chilensis*)* | VANCHI | | | 1 | 1 |
| Snowy Egret *(Egretta thula)* | EGRTHU | | | | 1 |
| Black-faced ibis (*Theristicus melanopis*)* | THEMEL | 1 | 1 | 1 | 1 |
| Andean condor *(Vultur gryphus)* | VULGRY | 1 | 1 | | |
| Black vulture *(Coragyps atratus)* | CORATR | | | | 1 |
| Turkey vulture *(Cathartes aura)* | CATAUR | 1 | | | |
| Cinereous harrier *(Circus cinereus)* | CIRCIN | 1 | | | |
| Chilean hawk *(Accipiter chilensis)* | ACCCHI | | | 1 | |
| Variable hawk (*Geranoaetus polyosoma*)* | GERPOL | 1 | 1 | 1 | 1 |
| White-throated hawk *(Buteo albigula)* | BUTALB | | | 1 | |
| Rufous-tailed hawk *(Buteo ventralis)* | BUTVEN | | | 1 | |
| Austral pygmy owl *(Glaucidium nana)* | GLANAN | | 1 | 1 | 1 |
| Striped woodpecker (*Veniliornis lignarius*)* | VENLIG | | 1 | 1 | 1 |
| Magellanic woodpecker (*Campephilus magellanicus*)* | CAMMAG | | 1 | 1 | 1 |
| Chilean flicker (*Colaptes pitius*)* | COLPIT | | 1 | 1 | 1 |
| Southern crested caracara (*Caracara plancus*)* | CARPLA | | 1 | 1 | 1 |
| Mountain caracara *(Phalcoboenus megalopterus)* | PHAMEG | 1 | 1 | | |
| Chimango caracara (*Milvago chimango*)* | MILCHI | 1 | 1 | 1 | 1 |
| American kestrel *(Falco sparverius)* | FALSPA | 1 | 1 | 1 | 1 |
| Aplomado falcon *(Falco femoralis)* | FALFEM | 1 | | | |
| Peregrine falcon *(Falco peregrinus)* | FALPER | | | 1 | 1 |
| Austral parakeet (*Enicognathus ferrugineus*)* | ENIFER | | 1 | 1 | 1 |
| Slender-billed parakeet *(Enicognathus leptorhynchus)* | ENILEP | | | 1 | |
| Black-throated huet-huet (*Pteroptochos tarnii*)* | PTETAR | | 1 | 1 | 1 |
| Chucao tapaculo (*Scelorchilus rubecula*)* | SCERUB | | 1 | 1 | 1 |
| Ochre-flanked tapaculo *(Eugralla paradoxa)* | EUGPAR | | | 1 | 1 |
| Magellanic tapaculo (*Scytalopus magellanicus*)* | SCYMAG | | 1 | 1 | 1 |
| Rufous-banded miner (*Geositta rufipennis*)* | GEORUF | 1 | 1 | | |

| | | | | | |
|---|---|---|---|---|---|
| White-throated treerunner (*Pygarrhichas albogularis*)* | PYGALB | | 1 | 1 | 1 |
| Patagonian forest earthcreeper (*Upucerthia saturatior*)* | UPUSAT | 1 | 1 | | |
| Buff-winged cinclodes (*Cinclodes fuscus*)* | CINFUS | 1 | 1 | 1 | |
| Grey-flanked cinclodes (*Cinclodes oustaleti*)* | CINOUS | 1 | 1 | | |
| Dark-bellied cinclodes (*Cinclodes patagonicus*)* | CINPAT | | | 1 | 1 |
| Thorn-tailed rayadito (*Aphrastura spinicauda*)* | APHSPI | | 1 | 1 | 1 |
| Des Murs`s wire-tail (*Sylviorthorhynchus desmursii*)* | SYLDES | | 1 | 1 | 1 |
| Plain-mantled tit-spinetail (*Leptasthenura aegithaloides*)* | LEPAEG | 1 | 1 | 1 | 1 |
| Sharp-billed canastero (*Asthenes pyrrholeuca*)* | ASTPYR | 1 | 1 | | |
| White-crested elaenia (*Elaenia albiceps*)* | ELAALB | 1 | 1 | 1 | 1 |
| Tufted tit-tyrant (*Anairetes parulus*)* | ANAPAR | | | 1 | 1 |
| Spot-billed ground-tyrant *(Muscisaxicola maculirostris)* | MUSMAU | 1 | | | |
| Ochre-naped Ground-tyrant *(Muscisaxicola flavinucha)* | MUSFLA | 1 | | | |
| Dark-faced ground-tyrant (*Muscisaxicola maclovianus*)* | MUSMAC | 1 | 1 | | |
| White-browed Ground-tyrant (*Muscisaxicola albilora*)* | MUSALB | 1 | 1 | | |
| Black-billed shrike-tyrant *(Agriornis montanus)* | AGRMON | 1 | | | |
| Great shrike-tyrant *(Agriornis lividus)* | AGRLIV | 1 | 1 | | |
| Fire-eyed diucon (*Xolmis pyrope*)* | XOLPYR | 1 | 1 | 1 | 1 |
| Patagonian tyrant (*Colorhamphus parvirostris*)* | COLPAR | | | 1 | 1 |
| Blue-and-white swallow (*Pygochelidon cyanoleuca*)* | PYGCYA | 1 | 1 | 1 | |
| Chilean swallow (*Tachycineta leucopyga*)* | TACMEY | 1 | 1 | 1 | 1 |
| Southern house wren (*Troglodytes musculus*)* | TROMUS | 1 | 1 | 1 | 1 |
| Austral thrush (*Turdus falcklandii*)* | TURFAL | 1 | 1 | 1 | 1 |
| Chilean mockingbird *(Mimus thenca)* | MIMTHE | | | | 1 |
| Greater yellow-finch *(Chirigue dorado)* | SICAUR | 1 | | | |
| Grassland yellow-finch (*Sicalis luteola*)* | SICLUT | | | | 1 |
| Patagonian sierra-finch (*Phrygilus patagonicus*)* | PHRPAT | 1 | 1 | 1 | 1 |
| Plumbeous sierra-finch (*Phrygilus unicolor*)* | PHRUNI | 1 | 1 | | |
| Yellow-bridled finch (*Melanodera xanthogramma*)* | MELXAN | 1 | 1 | | |
| Common diuca-finch (*Diuca diuca*)* | DIUDIU | 1 | 1 | | 1 |
| Rufous-collared Sparrow (*Zonotrichia capensis*)* | ZONCAP | 1 | 1 | 1 | 1 |
| Austral blackbird (*Curaeus curaeus*)* | CURCUR | 1 | 1 | 1 | 1 |
| Long-tailed meadowlark (*Sturnella loyca*)* | STULOY | | | | 1 |
| Black-chinned siskin *(Spinus barbatus)** | SPIBAR | 1 | 1 | 1 | 1 |

$^{\Delta}$ AL: Alpine, SA: Subalpine, OM: Old-growth montane forest, SM: Successional montane forest.
